# Cancer modeling by Transgene Electroporation in Adult Zebrafish (TEAZ)

**DOI:** 10.1101/297234

**Authors:** Scott J. Callahan, Stephanie Tepan, Yan M Zhang, Helen Lindsay, Alexa Burger, Nathaniel R. Campbell, Isabella S. Kim, Travis J. Hollmann, Lorenz Studer, Christian Mosimann, Richard M. White

**Author notes:** Corresponding Author: Richard White, M.D., Ph.D., 1275 York Avenue, MB 424, New York, N.Y. 10065.

## Abstract

Transgenic animals are invaluable for modeling cancer genomics, but often require complex crosses of multiple germline alleles to obtain the desired combinations. Zebrafish models have advantages in that transgenes can be rapidly tested by mosaic expression, but these typically lack spatial and temporal control of tumor onset, which limits their utility for the study of tumor progression and metastasis. To overcome these limitations, we have developed a method called Transgene Electroporation in Adult Zebrafish (TEAZ). TEAZ can deliver DNA constructs with promoter elements of interest to drive fluorophores, oncogenes, or CRISPR-Cas9-based mutagenic cassettes in specific cell types. Using TEAZ, we created a highly aggressive melanoma model by expression of BRAF^V600E^ in spatially constrained melanocytes in the context of p53 deficiency and Cas9-mediated inactivation of Rb1. Unlike prior models that take ~4 months to develop, we found that TEAZ leads to tumor onset in ~7 weeks and these develop in fully immunocompetent animals. As the resulting tumors initiated at highly defined locations, we could track their progression via fluorescence and documented deep invasion into tissues and metastatic deposits. TEAZ can be deployed to other tissues and cell types such as the heart with the use of suitable transgenic promoters. The versatility of TEAZ makes it widely accessible for rapid modeling of somatic gene alterations and cancer progression at a scale not achievable in other in vivo systems.

## Introduction

The zebrafish has become an increasingly applied model in cancer biology at the interface of basic discovery and preclinical animal experimentation. The high fecundity and relatively simple husbandry enable large experimental series *in vivo*. Early cancer models in zebrafish were largely developed using mutagens such as MNNG or DMBA, which were later supplanted by transgenic technologies(Beckwith et al., 2000; Pliss et al., 1982; Spitsbergen et al., 2000). The initial transgenic cancer models were developed by injecting 1-cell zebrafish embryos with DNA constructs containing a promoter and oncogene. For example, T-cell ALL was modeled by transgene driven expression of the *MYC* oncogene using the *rag2* promoter, and melanomas generated by expressing *BRAF*^*V600E*^ under the *mitfa* promoter in a *tp53*^−/-^ germline mutant background(Berghmans et al., 2005; Langenau et al., 2003; Patton et al., 2005).

Despite the documented power of transgenic tumor models for mechanism discovery and drug testing, current models have several significant drawbacks: i) the majority of established models do not exhibit spatio-temporal control, such that the timing and anatomical location of tumor onset remains variable(Kaufman et al., 2016; Patton et al., 2005; White et al., 2011), ii) they generally do not enable the introduction of serial somatic oncogenic events for modeling second and third hit mutations after onset(Mione and Trede, 2010; White et al., 2013); and iii) discerning multifocal primary tumors versus true metastatic spread of a single tumor is challenging(Kaufman et al., 2016). These issues all impose significant limitations for investigating tumor progression and metastasis.

Transplantation-based methods address some of these issues: tumors can be dissected off from a transgenic tumor-bearing animal or from patient-derived xenografts (PDXs), and then serially transplanted into recipient animals such as the *casper* recipient strain to allow detailed *in vivo* imaging(Fior et al., 2017; Heilmann et al., 2015; Hoffman, 2015; Siolas and Hannon, 2013; Tang et al., 2016; White et al., 2008; Zeng et al., 2017). Alternatively, stable cell lines can be generated from a transgenic animal, such as the ZMEL1 melanoma line, which can be similarly used for transplantation studies(Heilmann et al., 2015). While these transplantation approaches allow for precise spatiotemporal control and are amenable to imaging of metastasis, these experiments often require immunosuppression of the recipients either through irradiation or genetic manipulation of immune cells(Tang et al., 2016), in addition to the initial generation of the suitable cell line. Recent work from the Langenau lab has shown that syngeneic fish can be used as transplant recipients, but these require that the tumors be developed in that particular genetic background, somewhat limiting their broad use across cancer(Blackburn et al., 2011). Furthermore, transplantation cancer models implant foreign tumors into inherently artificial microenvironments.

A variety of Cre/Lox based approaches have been used in the zebrafish to control the cells that undergo initiation, including T-cell leukemia that can be controlled by mRNA injection(Langenau et al., 2005). A further modification uses CreERT2 drivers, such that both the cell type and timing of gene expression can be controlled(Hans et al., 2009), and inducible expression has also been achieved with heat shock Cre constructs(Hans et al., 2011). Despite these advances, there are still a paucity of verified Cre/Lox*-*based approaches to cancer in the zebrafish as the transgenic animals have proven time-consuming to create, and require complex breeding schemes to generate the final required genotypes.

Based on this, we wished to develop an approach that would enable introduction of oncogenic elements directly into adult somatic tissue in a spatio-temporally controlled manner. In this manuscript, we report oncogenesis via Transgene Electroporation into Adult Zebrafish (TEAZ), which models how tumors natively form in somatic tissues in a fully immunocompetent adult zebrafish. Electroporation applies electrical pulses to generate pores within the cell membrane, enabling extracellular biomolecules (including DNA) to enter the cell(Neumann et al., 1982; Wong and Neumann, 1982). Electroporation is widely used for stable introduction of DNA elements into cells in tissue culture and into chick and mouse embryos. Electroporation has occasionally been utilized in adult zebrafish but these studies have been limited to cell tracking and transient morpholino knockdowns and has never been applied to cancer modeling(Hoegler et al., 2011; Münch et al., 2013; Rambabu et al., 2005; Thummel et al., 2011). Several studies in mice have harnessed electroporation to introduce transgenes into select adult tissues, including retina, muscle, brain, and prostate(Maresch et al., 2016; Nomura et al., 2016; Swartz et al., 2001; Yarmush et al., 2014) and has been used to model tumors such as pancreatic cancer(Jung et al., 2014; Maresch et al., 2016; Park et al., 2014). However, these approaches require surgery of the mice and can only be limited to a small number of animals at a time, limiting the number of subjects that can be reasonably studied in each experiment. Based on these prior observations and the very large cohorts we can generate, we reasoned that direct electroporation of oncogenic transgenic constructs into the zebrafish would be a straightforward, highly scalable approach to model tumor formation in cells of interest. Because electrodes and DNA solutions can be placed at defined locations, TEAZ allows for delivery of multiple transgenes specifically to the anatomical locations of interest. We find that TEAZ allows for the development of complex, aggressive melanomas driven by expression of oncogenic BRAF^V600E^ in concert with loss of the tumor suppressors *p53* and *rb1*. These tumors are highly invasive and eventually metastasize to distant locations, unlike previous transgenic zebrafish melanoma models which do not generally metastasize(Patton et al., 2005). Given the wealth of functionally uncharted genetic lesions discovered from sequencing human tumors, TEAZ allows for testing of candidate mutations in a rapid, scalable *in vivo* system. More broadly, TEAZ can also be used to study somatic alteration of gene function in any adult tissue, which will have applications for diseases outside of cancer as well.

## Results

### Introduction of genetic elements into adult zebrafish via electroporation

The TEAZ method is designed to introduce genetic elements into specific locations within the adult zebrafish (Fig. 1a). The method has been optimized for the use of plasmids generated in *E. coli* and purified using standard plasmid purification protocols (midi preps). To test this protocol, we anesthetized adult zebrafish, and then injected 1.0 μl purified plasmid DNA (1000ng/μl) directly under the dorsal fin. The injected zebrafish was then quickly placed into an agarose mold to position the animal upright. Using paddle-shaped electrodes, we directed electrical pulses across the injected region (electroporator set to LV mode, 45V, 5 pulses, 60ms pulse length, 1s pulse interval). To maximize expression in the skin as opposed to deeper tissues, the cathode paddle can be placed just above the surface where the DNA was injected as this will pull the negatively charged DNA towards the surface adjacent to the injection site. After electroporation, we placed the anesthetized zebrafish into fresh water for recovery and maintenance through standard husbandry. Including anesthetization, DNA injection, and electroporation, the entire protocol takes approximately 45-60 seconds per animal.

**Fig. 1.**
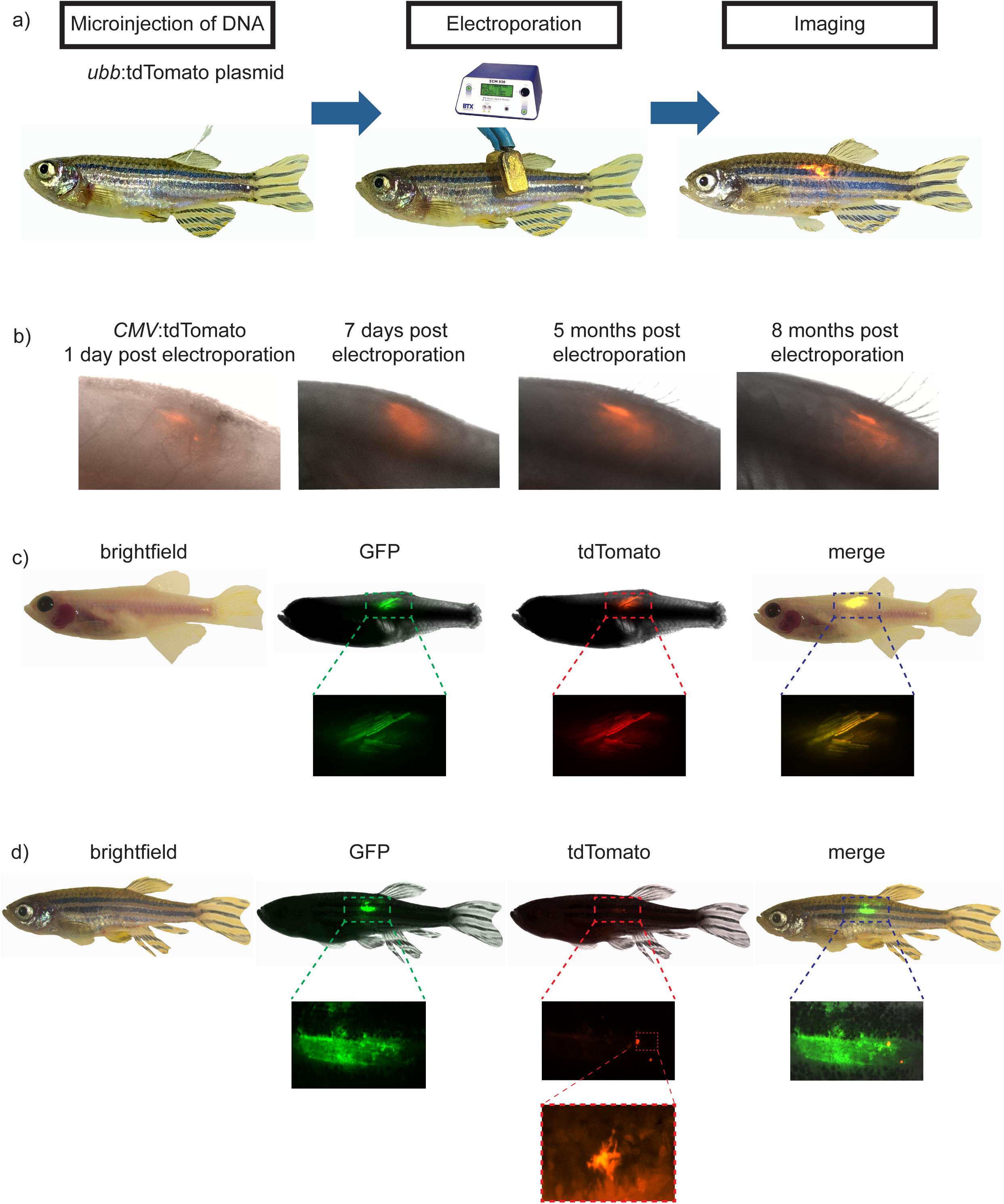
Transgene Electroporation into Adult Zebrafish (TEAZ). (a) Schematic representation of the method, applied for the introduction of *ubb*:tdTomato directly under the dorsal fin of adult zebrafish. The purified plasmid DNA (1μl of a 1000 ng/μl solution of *ubb*:dTomato) was injected into anesthetized zebrafish using a pulled glass micropipette. Electrical pulses are directed across the injected region (settings = LV mode, 45V, 5 pulses, 60ms pulse length, and 1s pulse interval). Reporter expression can be visualized by fluorescent microscopy (n=2/2). (b) Electroporation of a *CMV:tdTomato* plasmid was performed and the animal followed for a period of 8 months. The fluorescent signal can be visualized as early as 1 day post-electroporation (dpe), with intensity peaking around one week and maintaining for at least 8 months. (c) Multiple plasmids will co-integrate in TEAZ. *casper* zebrafish were electroporated with a total volume of 1.0 μl (0.5 μl of 1000ng/μl *ubb*:GFP and 0.5 μl of 1000 ng/μl *ubb*:tdTomato) and imaged using BF, GFP, and tdTomato (n=3/3), revealing co-expression of the plasmids. (d) Promoter specificity is maintained following TEAZ. *AB* fish were electroporated with 1.0 μl total volume (0.5 μl of 1000 ng/μl *ubb*:GFP and 0.5μl of 1000 ng/μl *mitfa*:tdTomato) and displayed highly restricted expression of the *mitfa* reporter plasmid, but widespread expression of the *ubb* plasmid (n=9/9). High-resolution imaging of the tdTomato-positive cells reveals a dendritic phenotype consistent with the melanocytic lineage.

To establish TEAZ, we injected wild-type zebrafish (*AB* strain) with a plasmid in which the zebrafish *ubiquitinB* promoter (*ubb*)(Mosimann et al., 2011) drives tdTomato expression (*ubb*:tdTomato). This vector was created using the Tol2 transposon system that is commonly deployed in the zebrafish(Balciunas et al., 2006; Kawakami, 2007; Kawakami et al., 2000; Kwan et al., 2007; Villefranc et al., 2007), although we did not utilize transposase mRNA for electroporation. After TEAZ, we imaged the electroporated zebrafish starting at 1 day post electroporation (1 dpe) and over the course of several months. Figure 1b shows an example animal, in which stable expression of tdTomato (under the *CMV* promoter) was detectable for up to 8 months. Similarly, the stable expression of *ubb*:GPF-p2a-tdTomato was also detectable after 8 months post electroporation (n = 4/4). We also tested whether the same *ubb* promoter and TEAZ would express in other parts of the animal by injecting it directly into the head. We observed robust expression of *ubb*:GFP in the head in 4 of the 5 animals electroporated (Fig. S1), indicating TEAZ-mediated transgene activity is not restricted to a particular location on the adult zebrafish body. At the site of electroporation there is initially a small area of tissue damage, that is rapidly healed within a week. We have not observed any systemic toxicity or obvious procedure-caused death in several hundred similarly electroporated animals.

### TEAZ allows for simultaneous expression of multiple plasmids

Electroporation of multiple plasmids into cultured cells *in vitro* generally leads to joint uptake of the plasmids by cells and subsequent co-expression of the transgenes. To test this in the zebrafish, we performed TEAZ with two plasmids in a single injection. We co-injected the *ubb*:tdTomato plasmid along with a *ubb*:GFP plasmid (each at 0.5 μl of 1000ng/μl plasmid stock) and then monitored fluorescence. High magnification views demonstrated that 100% of the transgene expressing cells were double positive for both GFP and tdTomato (n=3/3 fish) (Fig. 1c). Consequently, TEAZ can be expanded to express and combine multiple transgenes in the adult zebrafish skin.

### Maintenance of promoter specificity following electroporation

We next sought to determine whether TEAZ enables cell type-specific transgene expression. We co-electroporated *mitfa:tdTomato(Lister et al., 1999)* (which drives in melanocytes) and *ubb*:GFP (which drives ubiquitously) plasmids and then imaged the zebrafish using both tdTomato and GFP channels. A representative animal is shown in Figure 1d (n=9/9). As anticipated, we detected broad and strong expression of GFP from the *ubb* promoter(Mosimann et al., 2011). In contrast, we found highly limited expression of the *mitfa*:tdTomato plasmid. High resolution imaging of the tdTomato-positive cells revealed a dendritic appearance that is consistent with the appearance of mature melanocytes (Fig. 1d).

To test internal organs for TEAZ-mediated reporter delivery, we tested expression in the heart using the cardiomyocyte-specific *myl7* promoter driving GFP (*myl7:GFP*, formerly referred to as *cmlc*:GFP)(Huang et al., 2003). We injected and electroporated *myl7:GFP* plasmid directly into the beating heart muscle of an anesthetized adult zebrafish. We found strong and specific expression of GFP in the beating heart (n = 2/4) (Supplementary Figure 1c, Supplementary Movie 1). Importantly, when *myl7*:GFP was electroporated below the dorsal fin (n=5) and mitfa:tdTomato was electroporated into the heart (n=5) fluorescence was not detected. We conclude that TEAZ-mediated vector delivery maintains promoter specificity following electroporation, enabling us to target specific somatic cell types within specified regions of adult zebrafish.

### Melanoma initiation requires multiple transgenes

We next sought to apply TEAZ to directly model melanoma formation in adult zebrafish, circumventing embryonic manipulations. We and others have previously used a traditional germline melanoma transgenic in which the *mitfa* promoter drives oncogenic BRAF^V600E (Patton et al., 2005; White et al., 2013)^. In a *p53*^−/-^ deficient background, these transgenic animals develop a 100% penetrant melanoma at 4-12 months of age without any additional transgenes(Patton et al., 2005; White et al., 2011). This original transgenic was further extended using the *MiniCoopR* system(Ceol et al., 2011), in which the *mitfa*gene itself is knocked out, creating a strain with the genotype *mitfa*:BRAF^V600E^;*p53*^−/-^;*mitfa*^−/-^ (heretofore referred to as the “*triple*” strain). When this *triple* strain is injected at the 1-cell embryo stage with a “rescue” plasmid containing an *mitfa:mitfa* and *mitfa*:GFP cassette *in cis*, the resultant animals have rescued GFP+ melanocytes that all go on to develop GFP+ melanomas as adults(Ceol et al., 2011).

To test whether TEAZ is adaptable to this approach and could enable circumvention of initiating transgene expression at embryonic stages, we electroporated the *MiniCoopR:GFP* rescue cassette under the dorsal fin of *triple* strain adult zebrafish. We found that 8/10 injected animals developed GFP fluorescence at the site of injection, and remarkably, 1 of the animals developed rescued melanocytes. This indicates that it is possible to “rescue” melanophore development in a germline genetic defect (i.e. *mitfa*^−/-^) by directly electroporating a minigene cassette into adult somatic tissues. However, none of these animals went on to develop melanoma over a period of 4 months, a duration that leads to melanoma in embryo injection-based experiments. This observation suggests that in TEAZ, additional genetic hits are necessary above and beyond BRAF and *p53*^−/-^.

### TEAZ-mediated CRISPR of Rb1 stimulates melanoma in adults

In the *mitfa*^−/-^ mutant background, there are no mature melanocytes(Lister et al., 1999) since *mitfa* is required for expression of melanocytic genes such as *pmel* and *tyr*. We suspected that there might be melanocytic-precursor cells in the *mitfa*-BRAF^V600E^;*p53*^−/-^*;mitfa*^−/-^ background that are largely quiescent and not actively cycling. We therefore aimed to knock out the function of the tumor suppressor Rb1 using CRISPR-Cas9, since *p53* and *Rb1* mutations have a tendency to be concurrent in human melanoma patients as seen in the cBIO Portal (p=0.017)(Berger et al., 2012; Cerami et al., 2012; Gao et al., 2013; Hodis et al., 2012; Hugo et al., 2016; Krauthammer et al., 2012). Rb1 normally acts to keep cells arrested in G1, and we therefore reasoned that loss of its function might provoke the cells to proceed through the cell cycle and be more amenable to full malignant transformation(Yu et al., 2009).

To test this hypothesis, we employed the *triple* strain, and electroporated three plasmids: 1) *miniCoopR:GFP*, 2) *ubb:Cas9*, and 3) *zU6:sgRNA* against *rb1* (see Methods for details). We found that of the 9 electroporated animals, 8/9 developed rescued melanocytes and went on to establish aggressively growing GFP+ lesions with the phenotypic appearance of frank melanomas (Fig. 2). The tumors appeared within 3-7 weeks, in striking contrast to the 3-6 months typically required for standard embryo-injection transgenics. To confirm the effect was due to introduced *rb1* mutations, we dissected the dorsal fin (tumor) and tail fin (control normal tissue) from the same adult zebrafish for sequence analysis. Deep sequencing of the two fins and CrispRVariants-based allele analysis validated that the tumor contained two independent frameshift mutations in *rb1* at the PAM site that are characteristic of CRISPR mutations and that were not present in the control tail fin (Fig. S2) (Burger et al., 2016; Lindsay et al., 2016). These results reveal that TEAZ-mediated transformation of adult tissue can result in rapid melanoma onset using tumor-relevant genetic lesions To confirm that these lesions were truly tumorigenic, we followed a cohort of these TEAZ-treated zebrafish for a period of 4 months. By 5 weeks, primary melanomas could be visualized by both fluorescence and brightfield imaging. By 9 weeks, 4/8 remaining zebrafish had tumors that had rapidly progressed, and traversed the midline to the opposite side of the body. In 4/8 zebrafish, we noted evidence of GFP+ distant micrometastases in the head (Fig. 2c). To investigate further, we sacrificed 2 tumor-bearing animals along with 2 control animals and performed routine histology and anti-GFP staining. The first fish had a large protruding primary tumor with uniform GFP expression by fluorescence imaging. Histology of this tumor confirmed this with uniform anti-GFP staining, and H&E staining showed cells highly consistent with high-grade melanoma as determined by pathologist assessment (i.e. nuclear atypia and presence of melanin) (Fig. 3a). We noted extensive invasion into the muscle (Fig. 3b), which had not been previously seen in transgenic zebrafish melanoma modeling using BRAF^V600E^ with *p53*^−/-^. Along with this invasion phenotype, we identified micrometastases within the kidney (Fig. 3c) and attached to blood vessels (Fig. 3d). The second fish had an atypical tumor with variable GFP expression by fluorescence imaging, and histology showed a tumor of mixed histology in the vicinity of the injection needle: surface GFP-positive tumor cells consistent with a low grade melanoma, and a deeper tumor in the muscle that was GFP-negative consistent with a sarcoma. This second non-melanoma tumor is likely due to inactivation of both *rb1* and *p53* in the muscle, as we used ubiquitous:Cas9 in our studies and this combination is commonly found in sarcomas(Cerami et al., 2012; Gao et al., 2013; Gonin-Laurent et al., 2007; Pérot et al., 2010; Stratton et al., 1990). We did not find any GFP staining or abnormal H&E in either of the control animals (Fig. 3 and Supp Fig. 3c). Taken together, our findings demonstrate that the TEAZ method can rapidly and robustly give rise to tumors in a highly defined spatiotemporal manner. In contrast to previous transgenic zebrafish melanoma models, we also document evidence of progression and distant metastasis using TEAZ.

**Fig. 2.**
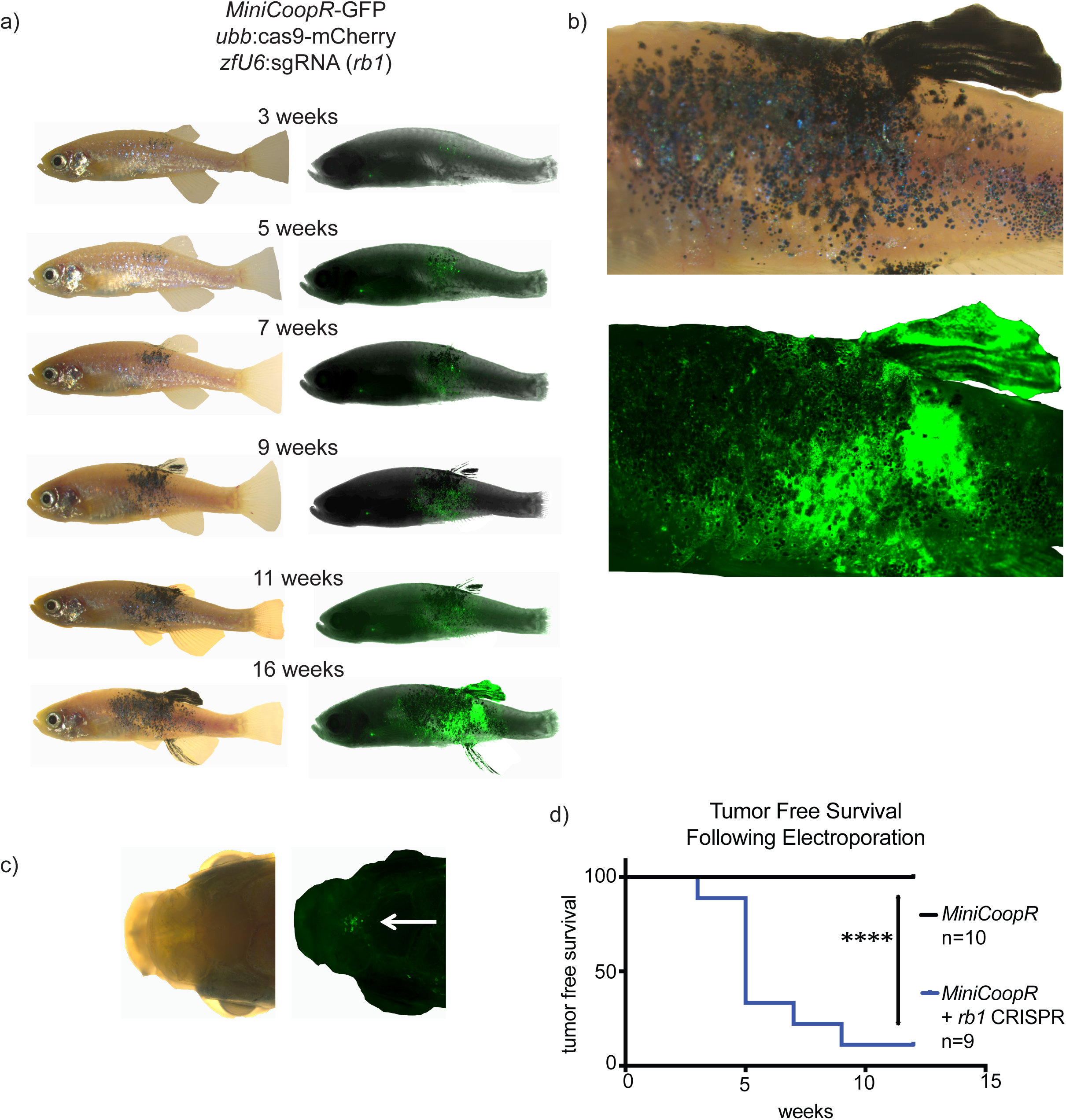
Generation of a novel melanoma model with TEAZ. (a) *mitfa*:BRAF^V600E^;*p53^−/-^*;*mitfa^−/-^* zebrafish (*triple* strain) were electroporated with the *miniCoopR:GFP* plasmid that both rescues melanocytes and expresses GFP under the *mitfa* promoter, with or without two additional plasmids to genetically knockout *rb1* (*ubb*:Cas9 and *zfU6*:sgRNA against *rb1*). The electroporated zebrafish were then imaged over time by both fluorescence and brightfield to monitor tumor development. Overall, 17/20 electroporated zebrafish had GFP+ cells. Tumor development in a representative zebrafish from the melanoma model including *rb1* knockout is shown. (b) Higher-magnification view of the tumor-bearing animal shown in (a) at 16 weeks post-electroporation. (c) At 9 weeks post-electroporation, 4/8 zebrafish had evidence of GFP+ distant micrometastases in the head. (d) The loss of *rb1* is essential for tumor initiation as visualized by the Kaplan-Meier curve comparing zebrafish electroporated with *miniCoopR:GFP* +/-*rb1 sgRNA*. Log-rank (Mantel-Cox) test p < 0.0001.

**Fig. 3.**
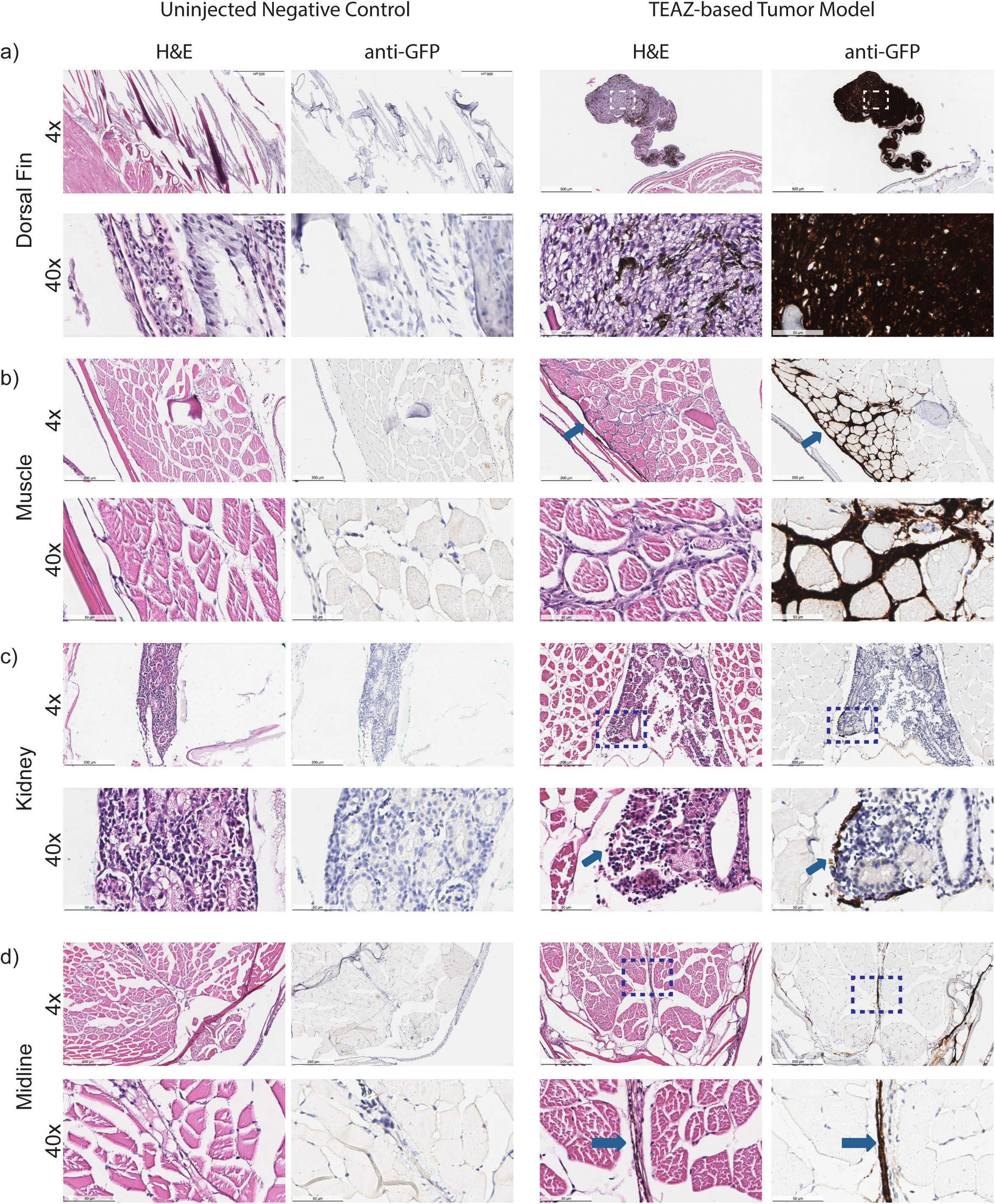
Melanoma model using TEAZ show evidence of rapid progression. (a) Pathology of tumor-bearing zebrafish (along with control zebrafish) at 16 weeks post-electroporation stained with hematoxylin/eosin or anti-GFP immunohistochemistry demonstrates a large primary tumor that is uniformly GFP-positive. (b) Histology reveals evidence of extensive invasion into the muscle (see arrow) along with micrometastatic sites within the (c) kidney or along (d) blood vessels (see arrows). Images are visualized at 4x and 40x where scale bars represent 500 μm and 50 μm respectively. Boxes represent the area enlarged at 40x.

### Somatic tumors are amenable to sequential transgenic manipulation

One of the major limitations of available genetic models is the ability to modify genes in a sequential order, mimicking *in vivo* tumor progression from malignant clones(McGranahan and Swanton, 2017). This limitation precludes the investigation into whether certain oncogenic events are driving initiation (which occur early) versus metastasis (when they may occur later). We therefore sought to determine if we could sequentially perform TEAZ to introduce new DNA elements into an already existing tumor. We selected a TEAZ-melanoma from the cohort above (4 months post-initial electroporation), and electroporated an *mitfa*:tdTomato plasmid directly into the tumor (Figure 4). Within 1 week after this second electroporation, we identified tdTomato-positive cells within the TEAZ-treated tumor (n=2/2). The tdTomato-cells have a dendritic appearance typical of a melanocytic cell. This indicates that existing tumors on adult zebrafish are amenable to TEAZ-mediated transgene delivery for sequential modeling of genetic lesions.

**Fig. 4.**
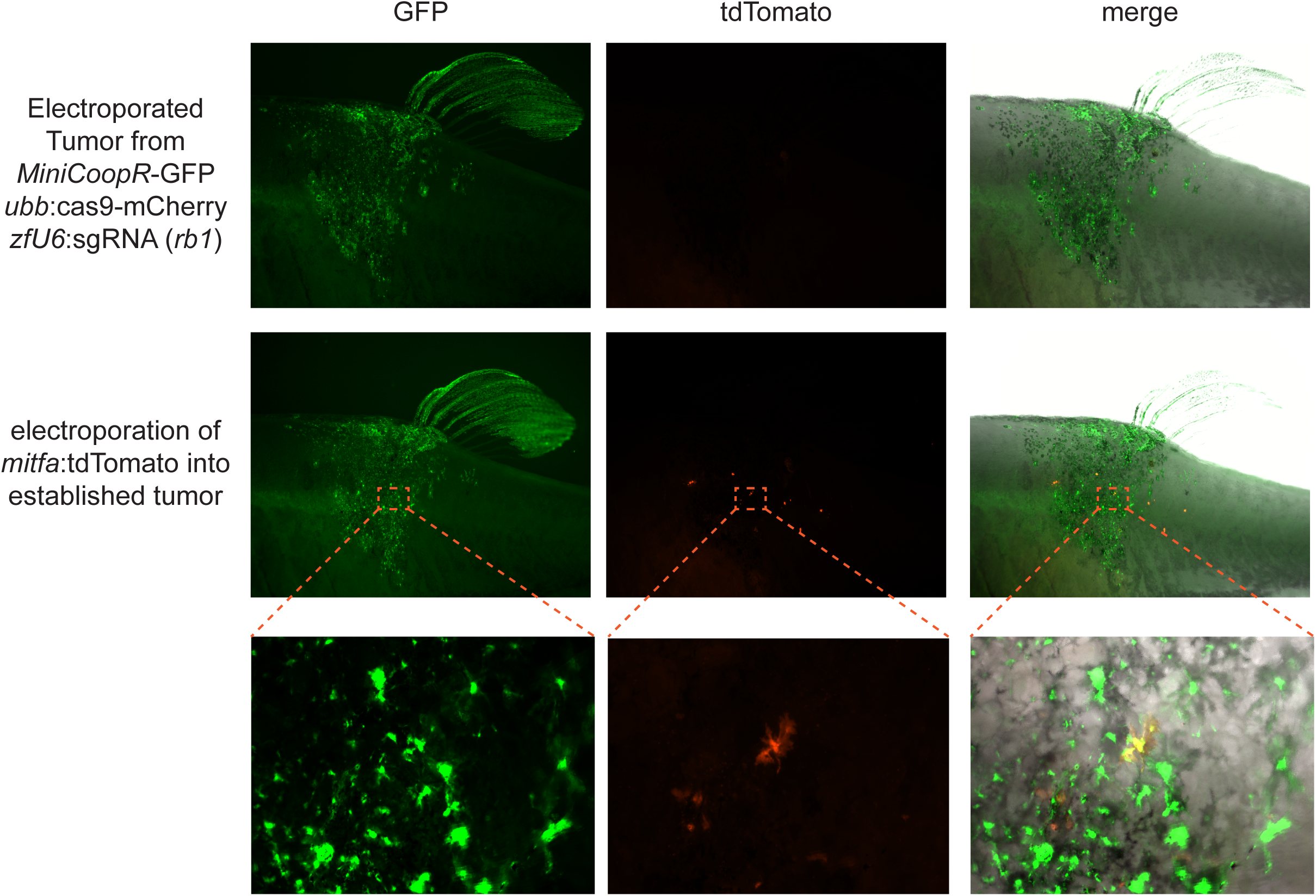
Cancer modeling with TEAZ enables sequential electroporation of transgenes. A tumor bearing fish (created with *rb1* sgRNA as in Figure 2) was imaged using GFP and tdTomato. As expected, only GFP+ tumor cells were seen with no expression in the tdTomato channel. This tumor was then electroporated with an *mitfa*:tdTomato plasmid and re-imaged 5 days later, showing areas that are now both GFP-negative and tdTomato-positive.

## Discussion

We have developed TEAZ, an electroporation-based approach for expressing transgenes and creating mutations in somatic cell types of interest within a region of interest in the adult zebrafish. We successfully applied TEAZ to the generation of malignant melanoma, and our results show TEAZ can be used for sequential electroporation to make increasingly complex tumor models. This model is amenable to initiating a tumor at a defined time and place, and will allow for a detailed analysis of tumor progression and metastasis in a fully immunocompetent zebrafish.

One major limitation of current transgenic cancer modeling in the zebrafish is the challenge of controlling both the timing and location of tumor initiation. Although both transplantation and Cre/Lox approaches can address some of these issues, neither fully solves the problem. Transplantation generally requires immune modulation of the recipient, and introduces relatively high cell burdens into tissue contexts that would not occur during natural tumor formation. Cre/Lox is extremely powerful but require multiple genetic crosses, and there are very few verified lines that exist for cancer modeling in the fish. In contrast, TEAZ rapidly allows for introduction of the required genetic elements in a multiplexed, complex manner.

Electroporation has been used as a mechanism for tumor initiation in mouse models of cancer. Both glioblastoma and oligoastrocytomas(Chen et al., 2016) have been induced into the developing fetus using *in utero* electroporation and the *piggyBAC* transposon system. In these studies, the transposase was included on the plasmid as a helper, although in our study we did not find Tol2 transposase to be necessary for highly stable transgene expression. Recent work(Jung et al., 2014; Maresch et al., 2016; Park et al., 2014) showed that plasmid delivery and electroporation could be used to initiate KRAS-driven pancreatic cancer in the adult mouse, and similar to our findings, can be complemented with CRISPR-Cas9-mediated genome editing. One exciting area that we believe TEAZ will open up is the possibility of new epithelial cancer models in the zebrafish, since that has not been developed on a very large scale.

One of the major advantages of our TEAZ system is the high efficiency we have observed. Once trained, we find that the rate of success of transgene expression approaches 100% of injected animals with 100% survival. In our study of the melanomas, we found that overall, 88% of the fish developed GFP+ cells by 3 weeks, and all of those fish subsequently went on to develop tumors by 7 weeks. This timing is an advance over the previous standard transgenic melanoma models, even with the more rapid *MiniCoopR* mosaic approaches that speed up tumors with the addition of oncogenes such as *SETDB1(Chen et al., 2016)*. In addition, embryo injection remains a relatively laborious process, whereas we find that the adult electroporation is simple and fast and can be easily taught to inexperienced users.

One possibility that our TEAZ system opens up is the ability to initiate adult-stage tumors in virtually any genetic background (i.e. the *casper* strain) or other existing transgenic line. In our studies using the miniCoopR background, we needed at least 3 genes to get efficient tumors: BRAF^V600E^, *p53*^−/-^ and *rb1*^−/-^. This may or may not be related to the specifics of the miniCoopR system as the melanocyte progenitors likely have to re-enter the cell cycle. We also noted that our tumors formed faster than typical miniCoopR tumors, even with the addition of genes such as SETDB1. This could relate to the specifics of the TEAZ approach or could simply be due to the accelerating effect of rb1 loss. It will be important for future studies to establish the minimum number of genetic elements necessary to drive tumor formation in wild-type backgrounds, which will allow TEAZ to be applied to existing transgenic lines that label a variety of interesting cell types such as T-cells(Dee et al., 2016), macrophages(Ellett et al., 2011) or endothelial cells(Lawson and Weinstein, 2002). Previously, creating transgenic tumors in a strain of interest required time-consuming breeding and genotyping to obtain the final genotype of interest. In contrast, with TEAZ, one can electroporate a combination of oncogenes and sgRNAs against tumor suppressors into any given genetic background directly, saving months of breeding and unused animals. To ensure specificity of the tumor types, it will be important to use tissue specific promoters to drive Cas9, to avoid mixed-histology tumors induced by ubiquitous Cas9 expression.

Metastasis and tumor progression have remained challenging to study using the zebrafish model; our results presented here suggest that TEAZ-mediated tumor modeling is amenable to studying metastasis in a high-throughput, immunocompetent model. Although transplantation-based approaches are powerful, they require immune system manipulation, such as irradiation or transplantation in genetically immuno-compromised zebrafish to counteract rejection(Heilmann et al., 2015; Tang et al., 2016; White et al., 2008; White et al., 2013). In contrast, TEAZ allows for tumor formation in fully immunocompetent animals. These features render the TEAZ model well-positioned to: (1) screen metastatic modulators to test rate, propensity, and latency; (2) selectively alter genes within specific cell types within the tumor microenvironment; (3) image the interplay between tumor cells and specific microenvironmental cell-types using widely available transgenic lines, and (4) introduce serial mutations to study order of progression or induce competition studies within a tumor.

## Methods

### Zebrafish Husbandry

Zebrafish were bred and maintained in the Zuckerman fish facility, in temperature (28°C), pH (7.4), and salinity-controlled conditions. All fish were maintained on a 14 hr on/10 hr off light cycle.

### Zebrafish Mutant Lines

Transgenic lines used in these studies included wild-type (*AB*), *casper(D’Agati et al., 2017; White et al., 2008)*, and the *triple* line(Ceol et al., 2011) (*mitfa*:BRAF^V600E^;*p53*^−/-^;*mitfa*^−/-^) (provided by the Houvras lab at Weill Cornell).

### Molecular Biology

Purified plasmids were generated using the Gateway system and isolated from *E.coli* using the Qiagen HiSpeed Plasmid Maxi Kit. The *MiniCoopR* vector was provided by the Houvras lab. The zebrafish *U6* promoter was cloned out of genomic DNA as previously described(Heilmann et al., 2015). The sgRNA against *rb1* was designed using the CHOPCHOP software and has the 20 bp sequence: GGCTCAGTGAGTTTGAACGG(Labun et al., 2016; Montague et al., 2014). The *ubb* promoter was as described previously(Mosimann et al., 2011), and the Cas9-mCherry fusion plasmid was subcloned from Addgene number *78313(Burger et al., 2016)*. All final plasmids were constructed using Gateway technology and the Tol2kit as previously described(Kawakami, 2007; Kwan et al., 2007).

List of all plasmids used: *ubb*:GFP(Mosimann et al., 2011) *and ubb*:tdTomato (both in Tol2kit plasmid backbone *#394*), *zfU6*:Rb1gRNA(Heilmann et al., 2015), *mitfa*:tdTomato(Lister et al., 2001), *myl7*:GFP(Huang et al., 2003), *MiniCoopR(Ceol et al., 2011)*, *ubb*:Cas9-mCherry fusion(Burger et al., 2016) (all in Tol2kit plasmid backbone *#395*).

### Electroporation of adult Zebrafish

At the time of electroporation, recipient adult zebrafish were anesthetized in 0.2% Tricaine. The plasmid of interest was resuspended at 1000ng/μl in ddH2O, and 1.0 μl was injected into the dorsal skin, brain or heart (using a pulled glass micropipette). No transposase mRNA was used in these studies. We have successfully tested a range of concentrations from 400-2000 ng/μl and from 0.5-2.0μl. Following injection, the zebrafish were immediately placed upright in an agarose mold for ease of handling and electrodes were placed on either side of the fish surrounding the injection site (Fig. 1a). The cathode paddle was generally placed on the same side as the injection to promote the DNA entering cells closer to the surface of the fish but the cathode and anode can be swapped to promote integration into cells deeper within the animal. We used the ECM 830 Electro Square Porator from BTX Harvard Apparatus and the Genepaddles, 3 x 5 mm. For all experiments described, the LV mode was used with a voltage of 45V, 5 pulses, 60ms pulse length, and 1s pulse interval. The electroporated zebrafish were immediately returned to flowing fresh water after electroporation. Electroporated zebrafish were imaged within 4 dpe to ensure successful TEAZ, and then serially imaged for up to 8 months using brightfield and fluorescence imaging.

### Imaging and image processing

Adult zebrafish were imaged using an upright Zeiss Discovery V16 equipped with a motorized stage, brightfield, and GFP and tdTomato filter sets. To acquire images, zebrafish were lightly anaesthetized with 0.2% Tricaine. Images were acquired with the Zeiss Zen software v1, and the post image processing was done using Fiji (Schindelin et al., 2012).

### Histology

Selected zebrafish were fixed in 4% PFA for 48 hours at 4°C and then paraffin embedded. Fish were sectioned at 5 uM, and then stained with H&E or anti-GFP. All histology was performed by Histowiz (http://www.histowiz.com) and reviewed by a pathologist (T.H.).

### Kaplan-Meier analysis

All animals were followed for up to 16 weeks and tumor free survival measured using the Kaplan-Meier method. The differences between the *miniCoopR* zebrafish with and without *rb1* knockout were analyzed using the log-rank statistics.

### MiSEQ analysis

Reads were mapped to the zebrafish genome version *GRCHz10* using bwa version 0.7.13-r1126. Mutation quantification was performed using CrispRVariants version 1.7.4(Lindsay et al., 2016).

### Author contributions

SJC, YMZ and RMW conceived the project. SJC, LS and RMW designed the experiments. SJC and ST performed the electroporation experiments. SJC, YMZ, IJK, NRC performed molecular cloning. HL, AB, and CM performed the CrispRVariants analysis. TJH performed histologic analysis. SJC and RMW wrote the manuscript.

## Acknowledgments

R.M.W. is supported by the NIH Director’s New Innovator Award (DP2CA186572), Mentored Clinical Scientist Research Career Development Award (K08AR055368), the Melanoma Research Alliance, The Starr Cancer Consortium, The Pershing Square Sohn Foundation, The Alan and Sandra Gerry Metastasis Research Initiative at the Memorial Sloan Kettering Cancer Center, The Harry J. Lloyd Foundation, Consano and and the Memorial Sloan Kettering Cancer Center Support Grant/Core Grant (P30CA008748). S.J.C is supported by the Kirschstein-NRSA predoctoral fellowship (F31) award as part of the National Cancer Institute of the National Institutes of Health under Award Number F31CA196305 as well as the Joanna M. Nicolay Melanoma Foundation Research Scholar Award 2014 and the Robert B. Catell Fellowship. N.R.C. is supported by a research grant from the Melanoma Research Foundation and by a Medical Scientist Training Program grant from the National Institute of General Medical Sciences of the National Institutes of Health under award number T32GM007739 to the Weill Cornell/Rockefeller/Sloan-Kettering Tri-Institutional MD-PhD Program. Y.M.Z. is supported by the Predoctoral to Postdoctoral Fellow Transition (F99/K00) Award as part of the National Cancer Institute of the National Institutes of Health under Award Number 1F99CA212436-01. C.M. received support through the Swiss National Science Foundation (SNSF) professorship (PP00P3_139093) and a Marie Curie Career Integration Grant from the European Commission (CIG PCIG14-GA-2013-631984) (C. Mosimann). A.B. and C.M. were further supported by the SwissBridge Foundation.

---

**Supplemental Fig. S1** TEAZ can be used to introduce transgenes in the adult brain and heart. (a) Purified plasmid encoding *ubb*:GFP (1μl of a 1000 ng/μl solution) was injected through the skull of an anesthetized zebrafish directly into the brain cavity using a pulled glass micropipette. The injected zebrafish is then electroporated across the dorsal-ventral axis of the head with the cathode positioned below the jaw and imaged for GFP fluorescence (n=4/5). (b) Pathology of the electroporated zebrafish with hematoxylin and eosin or immunochemistry against GFP to demonstrate reporter expression. Images are visualized at 4x and 20x where scale bars represent 500 μm and 100 μm respectively. (c) TEAZ can be extended to expression within the heart of adult zebrafish. *Casper* fish were injected into the heart through the gills with 1 μl of 1000ng/μl of a plasmid carrying the *myl7*:GFP transgene (along with a *ubb*:Cre cassette that is unrelated for the purposes of this study) (n=2/4). Video of the beating fluorescent heart can be seen in Supplementary Video 1.

**Supplemental Fig. S2.** A melanoma was induced as described in Main Figure 2, using *miniCoopR* and sgRNA against *rb1*. At 10 weeks post-electroporation, the melanoma was excised, along with control tissue from the tailfin of the same fish that was not electroporated. The tissues were digested for genomic DNA and the *rb1* locus was sequenced using multiplexed MiSEQ. In the tumor sample, we found a significant enrichment for two independent Cas9 induced mutations in *rb1* close to the *PAM* site (*-1:4D* and *-8:7D*) at an allele fraction over 2% each. A small number of these reads (0.04%) were found in the control tissue likely due to barcode contamination during sequencing multiplexing(Ballenghien et al., 2017). Yellow circle represents an inserted cytosine in -*1:1l,-3:9D*.

**Supplemental Fig. S3** Cancer model using TEAZ shows evidence of one tumor of mixed origin. (a) Fluorescent and brightfield imaging demonstrated a large primary tumor at the site of electroporation at 12 weeks post-electroporation with patchy GFP expression. (b) Imaging the zebrafish dorsally illustrates the large raised tumor from the contour of the animal. (c) Pathology of the tumor by hematoxylin and eosin or immunochemistry against GFP revealed that part of the deep tumor was GFP-negative and did not resemble a melanoma, but instead appeared consistent with a sarcoma. Images are visualized at 4x and 40x where scale bars represent 1 mm and 50 μm respectively. The blue boxes represent the area of sarcoma enlarged at 40x and the red boxes represent the area of melanoma enlarged at 40x.

**Supplemental Movie 1.** TEAZ can be used to introduce transgenes in the adult heart. *Casper* fish were injected into the heart through the gills with 1 μl of 1000ng/μl of a plasmid carrying the *myl7*:GFP transgene (along with a *ubb*:Cre cassette that is unrelated for the purposes of this study) (n=2/4). Video of the beating fluorescent heart was taken in the GFP channel.

## References

Balciunas, D., Wangensteen, K. J., Wilber, A., Bell, J., Geurts, A., Sivasubbu, S., Wang, X., Hackett, P. B., Largaespada, D. A., McIvor, R. S., et al. (2006). Harnessing a high cargo-capacity transposon for genetic applications in vertebrates. PLoS Genet. 2, e169.

Ballenghien, M., Faivre, N. and Galtier, N. (2017). Patterns of cross-contamination in a multispecies population genomic project: detection, quantification, impact, and solutions. BMC Biol. 15, 25.

Beckwith, L. G., Moore, J. L., Tsao-Wu, G. S., Harshbarger, J. C. and Cheng, K. C. (2000). Ethylnitrosourea induces neoplasia in zebrafish (Danio rerio). Lab. Invest. 80, 379–385.

Berger, M. F., Hodis, E., Heffernan, T. P., Deribe, Y. L., Lawrence, M. S., Protopopov, A., Ivanova, E., Watson, I. R., Nickerson, E., Ghosh, P., et al. (2012). Melanoma genome sequencing reveals frequent PREX2 mutations. Nature 485, 502–506.

Berghmans, S., Murphey, R. D., Wienholds, E., Neuberg, D., Kutok, J. L., Fletcher, C. D. M., Morris, J. P., Liu, T. X., Schulte-Merker, S., Kanki, J. P., et al. (2005). tp53 mutant zebrafish develop malignant peripheral nerve sheath tumors. Proc Natl Acad Sci USA 102, 407–412.

Blackburn, J. S., Liu, S. and Langenau, D. M. (2011). Quantifying the frequency of tumor-propagating cells using limiting dilution cell transplantation in syngeneic zebrafish. J. Vis. Exp. e2790.

Burger, A., Lindsay, H., Felker, A., Hess, C., Anders, C., Chiavacci, E., Zaugg, J., Weber, L. M., Catena, R., Jinek, M., et al. (2016). Maximizing mutagenesis with solubilized CRISPR-Cas9 ribonucleoprotein complexes. Development 143, 2025–2037.

Ceol, C. J., Houvras, Y., Jane-Valbuena, J., Bilodeau, S., Orlando, D. A., Battisti, V., Fritsch, L., Lin, W. M., Hollmann, T. J., Ferré, F., et al. (2011). The histone methyltransferase SETDB1 is recurrently amplified in melanoma and accelerates its onset. Nature 471, 513–517.

Cerami, E., Gao, J., Dogrusoz, U., Gross, B. E., Sumer, S. O., Aksoy, B. A., Jacobsen, A., Byrne, C. J., Heuer, M. L., Larsson, E., et al. (2012). The cBio cancer genomics portal: an open platform for exploring multidimensional cancer genomics data. Cancer Discov. 2, 401–404.

Chen, F., Becker, A. and LoTurco, J. (2016). Overview of transgenic glioblastoma and oligoastrocytoma CNS models and their utility in drug discovery. Curr. Protoc. Pharmacol. 72, 14.37.1-12.

Dee, C. T., Nagaraju, R. T., Athanasiadis, E. I., Gray, C., Fernandez Del Ama, L., Johnston, S. A., Secombes, C. J., Cvejic, A. and Hurlstone, A. F. L. (2016). CD4-Transgenic Zebrafish Reveal Tissue-Resident Th2− and Regulatory T Cell-like Populations and Diverse Mononuclear Phagocytes. J. Immunol. 197, 3520–3530.

D’Agati, G., Beltre, R., Sessa, A., Burger, A., Zhou, Y., Mosimann, C. and White, R. M. (2017). A defect in the mitochondrial protein Mpv17 underlies the transparent casper zebrafish. Dev. Biol. 430, 11–17.

Ellett, F., Pase, L., Hayman, J. W., Andrianopoulos, A. and Lieschke, G. J. (2011). mpeg1 promoter transgenes direct macrophage-lineage expression in zebrafish. Blood 117, e49–56.

Fior, R., Póvoa, V., Mendes, R. V., Carvalho, T., Gomes, A., Figueiredo, N. and Ferreira, M. G. (2017). Single-cell functional and chemosensitive profiling of combinatorial colorectal therapy in zebrafish xenografts. Proc Natl Acad Sci USA 114, E8234–E8243.

Gao, J., Aksoy, B. A., Dogrusoz, U., Dresdner, G., Gross, B., Sumer, S. O., Sun, Y., Jacobsen, A., Sinha, R., Larsson, E., et al. (2013). Integrative analysis of complex cancer genomics and clinical profiles using the cBioPortal. Sci. Signal. 6, pl1.

Gonin-Laurent, N., Hadj-Hamou, N. S., Vogt, N., Houdayer, C., Gauthiers-Villars, M., Dehainault, C., Sastre-Garau, X., Chevillard, S. and Malfoy, B. (2007). RB1 and TP53 pathways in radiation-induced sarcomas. Oncogene 26, 6106–6112.

Hans, S., Kaslin, J., Freudenreich, D. and Brand, M. (2009). Temporally-controlled site-specific recombination in zebrafish. PLoS ONE 4, e4640.

Hans, S., Freudenreich, D., Geffarth, M., Kaslin, J., Machate, A. and Brand, M. (2011). Generation of a non-leaky heat shock-inducible Cre line for conditional Cre/lox strategies in zebrafish. Dev. Dyn. 240, 108–115.

Heilmann, S., Ratnakumar, K., Langdon, E., Kansler, E., Kim, I., Campbell, N. R., Perry, E., McMahon, A., Kaufman, C., van Rooijen, E., et al. (2015). A quantitative system for studying metastasis using transparent zebrafish. Cancer Res. 75, 4272–4282.

Hodis, E., Watson, I. R., Kryukov, G. V., Arold, S. T., Imielinski, M., Theurillat, J.-P., Nickerson, E., Auclair, D., Li, L., Place, C., et al. (2012). A landscape of driver mutations in melanoma. Cell 150, 251–263.

Hoegler, K. J., Distel, M., Köster, R. W. and Horne, J. H. (2011). Targeting olfactory bulb neurons using combined in vivo electroporation and Gal4-based enhancer trap zebrafish lines. J. Vis. Exp.

Hoffman, R. M. (2015). Patient-derived orthotopic xenografts: better mimic of metastasis than subcutaneous xenografts. Nat. Rev. Cancer 15, 451–452.

Huang, C.-J., Tu, C.-T., Hsiao, C.-D., Hsieh, F.-J. and Tsai, H.-J. (2003). Germ-line transmission of a myocardium-specific GFP transgene reveals critical regulatory elements in the cardiac myosin light chain 2 promoter of zebrafish. Dev. Dyn. 228, 30–40.

Hugo, W., Zaretsky, J. M., Sun, L., Song, C., Moreno, B. H., Hu-Lieskovan, S., Berent-Maoz, B., Pang, J., Chmielowski, B., Cherry, G., et al. (2016). Genomic and Transcriptomic Features of Response to Anti-PD-1 Therapy in Metastatic Melanoma. Cell 165, 35–44.

Jung, S., Choi, H. J., Park, H. K., Jo, W., Jang, S., Ryu, J. E., Kim, W. J., Yu, E. S. and Son, W. C. (2014). Electroporation markedly improves Sleeping Beauty transposon-induced tumorigenesis in mice. Cancer Gene Ther. 21, 333–339.

Kaufman, C. K., Mosimann, C., Fan, Z. P., Yang, S., Thomas, A. J., Ablain, J., Tan, J. L., Fogley, R. D., van Rooijen, E., Hagedorn, E. J., et al. (2016). A zebrafish melanoma model reveals emergence of neural crest identity during melanoma initiation. Science 351, aad2197.

Kawakami, K. (2007). Tol2: a versatile gene transfer vector in vertebrates. Genome Biol. 8 Suppl 1, S7.

Kawakami, K., Shima, A. and Kawakami, N. (2000). Identification of a functional transposase of the Tol2 element, an Ac-like element from the Japanese medaka fish, and its transposition in the zebrafish germ lineage. Proc Natl Acad Sci USA 97, 11403–11408.

Krauthammer, M., Kong, Y., Ha, B. H., Evans, P., Bacchiocchi, A., McCusker, J. P., Cheng, E., Davis, M. J., Goh, G., Choi, M., et al. (2012). Exome sequencing identifies recurrent somatic RAC1 mutations in melanoma. Nat. Genet. 44, 1006–1014.

Kwan, K. M., Fujimoto, E., Grabher, C., Mangum, B. D., Hardy, M. E., Campbell, D. S., Parant, J. M., Yost, H. J., Kanki, J. P. and Chien, C.-B. (2007). The Tol2kit: a multisite gateway-based construction kit for Tol2 transposon transgenesis constructs. Dev. Dyn. 236, 3088–3099.

Labun, K., Montague, T. G., Gagnon, J. A., Thyme, S. B. and Valen, E. (2016). CHOPCHOP v2: a web tool for the next generation of CRISPR genome engineering. Nucleic Acids Res. 44, W272–6.

Langenau, D. M., Traver, D., Ferrando, A. A., Kutok, J. L., Aster, J. C., Kanki, J. P., Lin, S., Prochownik, E., Trede, N. S., Zon, L. I., et al. (2003). Myc-induced T cell leukemia in transgenic zebrafish. Science 299, 887–890.

Langenau, D. M., Feng, H., Berghmans, S., Kanki, J. P., Kutok, J. L. and Look, A. T. (2005). Cre/lox-regulated transgenic zebrafish model with conditional myc-induced T cell acute lymphoblastic leukemia. Proc Natl Acad Sci USA 102, 6068–6073.

Lawson, N. D. and Weinstein, B. M. (2002). In vivo imaging of embryonic vascular development using transgenic zebrafish. Dev. Biol. 248, 307–318.

Lindsay, H., Burger, A., Biyong, B., Felker, A., Hess, C., Zaugg, J., Chiavacci, E., Anders, C., Jinek, M., Mosimann, C., et al. (2016). CrispRVariants charts the mutation spectrum of genome engineering experiments. Nat. Biotechnol. 34, 701–702.

Lister, J. A., Robertson, C. P., Lepage, T., Johnson, S. L. and Raible, D. W. (1999). nacre encodes a zebrafish microphthalmia-related protein that regulates neural-crest-derived pigment cell fate. Development 126, 3757–3767.

Lister, J. A., Close, J. and Raible, D. W. (2001). Duplicate mitf genes in zebrafish: complementary expression and conservation of melanogenic potential. Dev. Biol. 237, 333–344.

Maresch, R., Mueller, S., Veltkamp, C., Öllinger, R., Friedrich, M., Heid, I., Steiger, K., Weber, J., Engleitner, T., Barenboim, M., et al. (2016). Multiplexed pancreatic genome engineering and cancer induction by transfection-based CRISPR/Cas9 delivery in mice. Nat. Commun. 7, 10770.

McGranahan, N. and Swanton, C. (2017). Clonal heterogeneity and tumor evolution: past, present, and the future. Cell 168, 613–628.

Mione, M. C. and Trede, N. S. (2010). The zebrafish as a model for cancer. Dis. Model. Mech. 3, 517–523.

Montague, T. G., Cruz, J. M., Gagnon, J. A., Church, G. M. and Valen, E. (2014). CHOPCHOP: a CRISPR/Cas9 and TALEN web tool for genome editing. Nucleic Acids Res. 42, W401–7.

Mosimann, C., Kaufman, C. K., Li, P., Pugach, E. K., Tamplin, O. J. and Zon, L. I. (2011). Ubiquitous transgene expression and Cre-based recombination driven by the ubiquitin promoter in zebrafish. Development 138, 169–177.

Münch, J., González-Rajal, A. and de la Pompa, J. L. (2013). Notch regulates blastema proliferation and prevents differentiation during adult zebrafish fin regeneration. Development 140, 1402–1411.

Neumann, E., Schaefer-Ridder, M., Wang, Y. and Hofschneider, P. H. (1982). Gene transfer into mouse lyoma cells by electroporation in high electric fields. EMBO J. 1, 841–845.

Nomura, T., Nishimura, Y., Gotoh, H. and Ono, K. (2016). Rapid and efficient gene delivery into the adult mouse brain via focal electroporation. Sci. Rep. 6, 29817.

Park, J.-S., Lim, K.-M., Park, S. G., Jung, S. Y., Choi, H.-J., Lee, D. H., Kim, W.-J., Hong, S.-M., Yu, E.-S. and Son, W.-C. (2014). Pancreatic cancer induced by in vivo electroporation-enhanced sleeping beauty transposon gene delivery system in mouse. Pancreas 43, 614–618.

Patton, E. E., Widlund, H. R., Kutok, J. L., Kopani, K. R., Amatruda, J. F., Murphey, R. D., Berghmans, S., Mayhall, E. A., Traver, D., Fletcher, C. D. M., et al. (2005). BRAF mutations are sufficient to promote nevi formation and cooperate with p53 in the genesis of melanoma. Curr. Biol. 15, 249–254.

Pérot, G., Chibon, F., Montero, A., Lagarde, P., de Thé, H., Terrier, P., Guillou, L., Ranchère, D., Coindre, J.-M. and Aurias, A. (2010). Constant p53 pathway inactivation in a large series of soft tissue sarcomas with complex genetics. Am. J. Pathol. 177, 2080–2090.

Pliss, G. B., Zabezhinski, M. A., Petrov, A. S. and Khudoley, V. V. (1982). Peculiarities of N-nitramines carcinogenic action. Arch. Geschwulstforsch. 52, 629–634.

Rambabu, K. M., Rao, S. H. N. and Rao, N. M. (2005). Efficient expression of transgenes in adult zebrafish by electroporation. BMC Biotechnol. 5, 29.

Schindelin, J., Arganda-Carreras, I., Frise, E., Kaynig, V., Longair, M., Pietzsch, T., Preibisch, S., Rueden, C., Saalfeld, S., Schmid, B., et al. (2012). Fiji: an open-source platform for biological-image analysis. Nat. Methods 9, 676–682.

Siolas, D. and Hannon, G. J. (2013). Patient-derived tumor xenografts: transforming clinical samples into mouse models. Cancer Res. 73, 5315–5319.

Spitsbergen, J. M., Tsai, H. W., Reddy, A., Miller, T., Arbogast, D., Hendricks, J. D. and Bailey, G. S. (2000). Neoplasia in zebrafish (Danio rerio) treated with 7,12-dimethylbenz[a]anthracene by two exposure routes at different developmental stages. Toxicol. Pathol. 28, 705–715.

Stratton, M. R., Moss, S., Warren, W., Patterson, H., Clark, J., Fisher, C., Fletcher, C. D., Ball, A., Thomas, M. and Gusterson, B. A. (1990). Mutation of the p53 gene in human soft tissue sarcomas: association with abnormalities of the RB1 gene. Oncogene 5, 1297–1301.

Swartz, M., Eberhart, J., Mastick, G. S. and Krull, C. E. (2001). Sparking new frontiers: using in vivo electroporation for genetic manipulations. Dev. Biol. 233, 13–21.

Tang, Q., Moore, J. C., Ignatius, M. S., Tenente, I. M., Hayes, M. N., Garcia, E. G., Torres Yordán, N., Bourque, C., He, S., Blackburn, J. S., et al. (2016). Imaging tumour cell heterogeneity following cell transplantation into optically clear immune-deficient zebrafish. Nat. Commun. 7, 10358.

Thummel, R., Bailey, T. J. and Hyde, D. R. (2011). In vivo electroporation of morpholinos into the adult zebrafish retina. J. Vis. Exp. e3603.

Villefranc, J. A., Amigo, J. and Lawson, N. D. (2007). Gateway compatible vectors for analysis of gene function in the zebrafish. Dev. Dyn. 236, 3077–3087.

White, R. M., Sessa, A., Burke, C., Bowman, T., LeBlanc, J., Ceol, C., Bourque, C., Dovey, M., Goessling, W., Burns, C. E., et al. (2008). Transparent adult zebrafish as a tool for in vivo transplantation analysis. Cell Stem Cell 2, 183–189.

White, R. M., Cech, J., Ratanasirintrawoot, S., Lin, C. Y., Rahl, P. B., Burke, C. J., Langdon, E., Tomlinson, M. L., Mosher, J., Kaufman, C., et al. (2011). DHODH modulates transcriptional elongation in the neural crest and melanoma. Nature 471, 518–522.

White, R., Rose, K. and Zon, L. (2013). Zebrafish cancer: the state of the art and the path forward. Nat. Rev. Cancer 13, 624–636.

Wong, T. K. and Neumann, E. (1982). Electric field mediated gene transfer. Biochem. Biophys. Res. Commun. 107, 584–587.

Yarmush, M. L., Golberg, A., Serša, G., Kotnik, T. and Miklavčič, D. (2014). Electroporation-based technologies for medicine: principles, applications, and challenges. Annu. Rev. Biomed. Eng. 16, 295–320.

Yu, H., McDaid, R., Lee, J., Possik, P., Li, L., Kumar, S. M., Elder, D. E., Van Belle, P., Gimotty, P., Guerra, M., et al. (2009). The role of BRAF mutation and p53 inactivation during transformation of a subpopulation of primary human melanocytes. Am. J. Pathol. 174, 2367–2377.

Zeng, A., Ye, T., Cao, D., Huang, X., Yang, Y., Chen, X., Xie, Y., Yao, S. and Zhao, C. (2017). Identify a Blood-Brain Barrier Penetrating Drug-TNB using Zebrafish Orthotopic Glioblastoma Xenograft Model. Sci. Rep. 7, 14372.

